# Quantitative mapping of cerebral oxygen metabolism using breath-hold calibrated fMRI

**DOI:** 10.1101/2021.04.08.438939

**Authors:** M Germuska, RC Stickland, AM Chiarelli, HL Chandler, RG Wise

**Affiliations:** Cardiff University; Northwestern University; Department of Neuroscience, Imaging and Clinical Sciences, Institute for Advanced Biomedical Technologies, University G. D Annunzio of Chieti and Pescara, Via Luigi Polacchi 13, 66100 Chieti, Italy

## Abstract

Magnetic resonance imaging (MRI) offers the possibility to non-invasively map the rate of cerebral metabolic oxygen consumption (CMRO_2_), which is essential for understanding and monitoring neural function in both health and disease. Existing methods of mapping CMRO_2_, based on respiratory modulation of arterial spin labelling (ASL) and blood oxygen level dependent (BOLD) signals, require lengthy acquisitions and independent modulation of both arterial oxygen and carbon dioxide levels. Here, we present a new simplified method for mapping the rate of cerebral oxygen metabolism that can be performed using a simple breath-holding paradigm. The method incorporates flow-diffusion modelling of oxygen transport and physiological constraints to create a non-linear mapping between the maximum BOLD signal, M, baseline blood flow (CBF_0_), and CMRO_2_. A gradient boosted decision tree is used to learn this mapping directly from simulated MRI data. Modelling studies demonstrate that the proposed method is robust to variation in cerebral physiology and metabolism. This new gas-free methodology offers a rapid and pragmatic alternative to existing dual-calibrated methods, removing the need for specialist respiratory equipment and long acquisition times. In-vivo testing of the method, using an 8-minute 45 second protocol of repeated breath-holding, was performed on 15 healthy volunteers, producing quantitative maps of cerebral blood flow (CBF), oxygen extraction fraction (OEF), and CMRO_2_.

## 1 Introduction

A continuous supply of oxygen to the brain is essential for life and any restriction of this supply can have significant consequences for individuals ^1^. The current gold standard for mapping the cerebral metabolic rate of oxygen consumption (CMRO_2_) is 15-oxygen labelled positron emission tomography ^2^. However, this method has substantial drawbacks, including long acquisition times, the use of ionizing radiation, and the need for local production of 15-oxygen labelled tracers. Due to these limitations, there is great interest in developing rapid and non-invasive methods that can be used to safely and quickly map CMRO_2_. In recent years, a number of MRI methods ^3–6^ has been proposed as an alternative to PET based methods. One promising approach is the so-called dual-calibrated fMRI method ^7–9^. This approach combines biophysical modelling of the blood oxygen level dependent (BOLD) signal, and modulation of cerebral blood flow and blood oxygenation with controlled respiratory stimuli. This approach has been successfully applied in disease ^10, 11^, sports related head impact ^12^, and pharmacological modulation studies ^13^. However, the dual-calibrated method requires specialist respiratory equipment and experienced operators to acquire the data. These requirements limit the wider adoption of such a method both in research and clinical settings. In this work we present a straightforward gas-free approach for estimating the resting cerebral metabolic rate of oxygen consumption (CMRO_2,0_). The method uses a simple repeated breath-holding paradigm to provide robust estimates of CMRO_2,0_. The data can be acquired in less than 10 minutes with minimal operator training.

The dual-calibrated method can be conceptualised as making two independent measurements of M (the maximum possible BOLD signal); one measurement is dependent on the oxygen extraction fraction (OEF), and one is not ^7^. OEF is then estimated by assigning a value that brings the two measurements of M into agreement. In our proposed method only one measurement of M is required. Estimates of M can be acquired with a calibrated-fMRI experiment using either a hypercapnic gas challenge ^14^ or a breath-holding protocol ^15^, or by transforming a measurement of R_2_’ ^16^. In the work presented here we have implemented the modelling and analysis for a repeated breath-holding protocol, allowing mapping of CMRO_2,0_ from a simple imaging protocol without the need for administration of modified gas mixtures.

The proposed method is based on a new formulation of M, which includes a simple model of oxygen exchange ^17–19^, and substitutes the deoxyhaemoglobin sensitive blood volume (CBV_v_) for an appropriately scaled capillary blood volume (CBV_cap_). By combining the newly formulated expression of M with reasonable physiological constraints, we can create a non-linear mapping from M and the resting cerebral blood flow (CBF_0_) to CMRO_2,0_. For in-vivo data analysis we employ a machine learning methodology that estimates CMRO_2,0_ directly from the MRI data, without estimating the intermediate parameters. We have previously shown that such an approach, trained on simulated data, is able to make robust estimates of physiological and metabolic parameters when applied to the dual-calibrated fMRI method ^19^.

We adopted the same concept as used in our recent publication ^19^ and estimate CMRO_2,0_ from a Fourier transformed representation of the MRI time series data. In the proposed implementation we use a popular gradient boosted decision tree algorithm, LightGBM^20^, to learn the mapping between the simulated MRI data and CMRO_2,0_. The method is tested via further simulations and in-vivo data acquired from with 15 healthy volunteers.

## 2 MRI Data Modelling

Modelling of the steady-state BOLD signal shows that the rate of transverse relaxation due to deoxyhaemoglobin can be expressed as shown in equation 1 ^14, 21^.

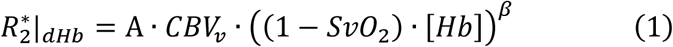

Where 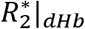 is the transverse relaxation rate due to deoxyhaemoglobin. CBV_v_ is the fractional BOLD sensitive blood volume, which is composed of the venous blood volumes, and capillary blood volumes that are exchanging oxygen with the tissue. β is a field strength dependent constant, and A is a proportionality constant related to the field strength and vessel geometry (in units of s^−1^g-^β^dL^β^). [Hb] is the haemoglobin concentration (g/dL) and SvO_2_ is the draining venous oxygen saturation.

The maximum BOLD signal, M, is obtained simply by multiplying 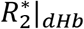 by the acquisition echo time, TE ^14^.

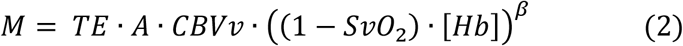

Using this definition of M, the generic BOLD signal model that describes the signal change due to modulation of arterial oxygen content (CaO_2_), CBF, or CMRO_2_ can be expressed as shown in equation 3a ^19^.

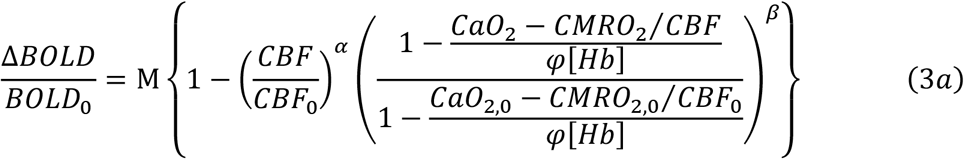

Where, Δ*BOLD/BOLD*_0_ is the fractional change in BOLD signal, φ is the oxygen binding capacity of haemoglobin (1.34 ml/g), and the subscript 0 denotes the resting value.

In agreement with the previous literature, we assume that brief periods of breath-holding are iso-metabolic ^15, 22, 23^. Additionally, we follow recent QSM modelling ^24^ and vascular measurements ^25^ in setting the Grubb exponent, α, to zero. Thus, for a breath-holding stimulus, the generalised BOLD model of equation 3a is simplified to equation 3b. Unlike the standard hypercapnic calibration equation ^14^, we include the arterial oxygen content (CaO_2_) as this is known to change during breath-holding ^26^. Because of this, the equation also maintains a dependence on CMRO_2,0_.

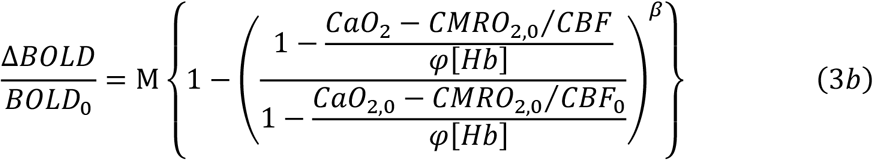

Typically, the influence of CaO_2_ on the BOLD signal is ignored in calibration experiments based on breath-holding ^15, 22, 27^. However, here it is included for a more complete description of the BOLD signal. Upon completion of a breath-holding experiment equation 3b will have two unknowns, M and CMRO_2,0_. Thus, it initially appears that we cannot use a breath-holding paradigm to solve for either parameter. A similar problem is solved in the dual-calibrated methodology by including a hyperoxic measurement, which is predominately sensitive to CBV_v_ ^28^. Here we resolve the problem by employing oxygen diffusion modelling and physiological constraints during fitting.

Following the modelling of ^17, 18, 29^ we can express the resting cerebral metabolic rate of oxygen metabolism as shown in equation 4a.

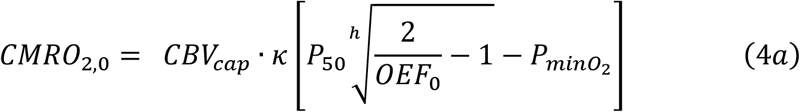

Where CBV_cap_ is the capillary blood volume that exchanges oxygen with the tissue, κ is the effective permeability of capillary endothelium and brain tissue (μmol/mmHg/ml/min), P50 is the blood oxygen tension at which haemoglobin is 50% saturated, h is the Hill coefficient (equal to 2.8) and PminO_2_ is the minimum oxygen tension at the mitochondria. In the modelling we calculate the value of P50 from a measure of end-tidal carbon dioxide tension and we assume a fixed value for κ of 3 μmol/mmHg/ml/min ^19^.

The capillary blood volume exchanging oxygen with the tissue, CBV_cap_, is a fraction of the BOLD sensitive blood volume (CBV_v_), i.e. *CBV*_*ν*_ = *ρ* · *CBV*_*cap*_. Thus, by re-arranging equation 4a in terms of CBV_v_ (equation 4b) and substituting into the definition of M we can derive an alternative definition of M as expressed by equation 5.

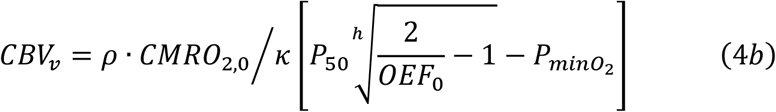

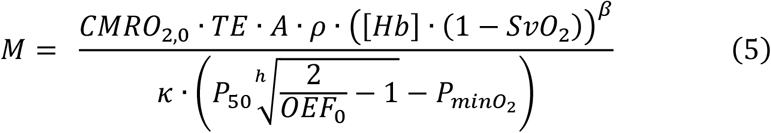

This definition of M effectively replaces the unknown parameter CBV_v_ with two measurable parameters, CaO_2,0_ and CBF_0_ and two unknown parameters, ρ and PminO_2_. This can be seen by examining equation 5 and expanding CMRO_2,0_ via the Fick principle, *CMRO*_2,0_ = *CaO*_2,0_ · *CBF*_0_ · *OEF*_0_. The standard definition of M contains TE, A, [Hb], and SvO_2_, thus there is already an implicit dependence on OEF_0_ (assuming arterial blood is close to, or fully saturated). The Hill coefficient and the effective permeability of brain tissue can be assumed constant, and the value of P50 can be estimated from the end-tidal carbon dioxide tension. Thus, the remaining unknown parameters are ρ and PminO_2_.

By definition PminO_2_ must lie between 0 mmHg and the oxygen tension of the capillary bed, PcapO_2_. In vivo studies suggest the oxygen tension at the brain mitochondria is low in healthy brain ^17, 30^, with an average value of approximately 8.5 mmHg. In this work we use a gamma distribution to model PmitO_2_, maintaining the same median value but allowing variation between zero and PcapO_2_. Thus, including the entire range of feasible values for the mitochondrial oxygen tension in the healthy and diseased brain.

The in-vivo variation in ρ has not been studied directly, however, we can gain some insight by considering the variation in vascular volume fractions with ageing and disease. For example, studies of capillary density in the human brain report age related decreases of approximately 16%, while studies in rat suggest that such vascular changes are mirrored by alterations in venular blood volume, with an approximate 20% decrease in venular blood observed across the life span of rats ^31^. Large reductions in vascular density have been observed in aging patients with neurodegeneration. For example, Buee et al. ^32^ reported a 38% decrease in vessel density in elderly AD patients compared to a healthy 49-year-old. However, the fractional change in the capillary density is reported to be of a similar magnitude, suggesting minimal alterations in ρ ^33, 34^.

MRI studies of stroke using vessel size imaging (VSI) report an increase in the mean vessel size and a reduction in vessel density. Although histological studies confirm a significant reduction in vessel density, they do not report any change in the mean vessel size ^35^. The in-vivo observation of an increase in vessel size is likely to be due to vasodilation of arterioles, and thus the capillary fraction of the BOLD sensitive vasculature is likely to remain unchanged even in stroke.

Taken together these findings suggest there is little variation in the relationship between capillary blood volume and the BOLD sensitive blood volume in a wide array of physiological and pathological conditions including ageing, neurodegeneration, and stroke. In our modelling we chose the range of ρ to be 2 to 3.33, giving a capillary blood volume of 20 to 40% of total blood volume, when the arterial contribution is assumed to be 20 to 30% ^36^. This range was chosen to offer reasonable physiological variation around a typical capillary volume of 33% ^37, 38^.

The scaling factor, A, in the BOLD equation is similar to β, in that they both capture information related to vessel geometry and field strength ^39^. While a fixed value is normally given to β, the value of A is normally left undefined. However, we are able to approximate A if we assume that the majority of the BOLD signal arises from the extravascular space around veins and 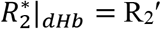 This simplification follows that used by Blockley et al. who used R_2_′ as a surrogate for M in calibrated fMRI studies. ^16, 40^. Blockley et al demonstrate that R_2_′ captures most of the BOLD signal variation and predict a tight correlation between R_2_′ and M. Thus, with reference to equation 1, a reasonable approximation of A can be obtained from in-vivo measurements of R_2_′, CBV_v_, and SvO_2_. The cortical R_2_′ is approximately is 3 s^−1^ at 3T ^41^, while the BOLD sensitive blood volume has been estimated to be between 1.75% ^6^ and 3.6% ^42^. Assuming an average [Hb] of 14 g/dL, an SvO_2_ of 0.6, and taking the mean CBV_v_ of 2.5% we obtain a value of 14 s^−1^g-^β^dL^β^ for A at 3T.

By taking this estimate of A as the mean value and including variance to capture the physiological uncertainty, we are able to perform Monte-Carlo simulations to explore the relationship between M, CBF_0_ and OEF_0_. We performed these simulations with both the standard definition of M and the newly proposed definition (equation 5). Table 1 outlines the physiological parameters and values used in the Monte-Carlo simulations.

**Table 1.** Parameter ranges and distributions (Γ(shape, rate), *N*(mean, variance)) for two sets of Monte-Carlo simulations (**a** and **b**). The simulations model the relationship between CBF_0_, M, and OEF_0_ and are summarised in figure 1.

| Parameter Description | Symbol | Simulation<br>a | Simulation<br>b |
| --- | --- | --- | --- |
| Maximum BOLD signal | M | 0.001 – 0.4 | 0.001 – 0.4 |
| Model includes oxygen diffusion | - | × | ✓ |
| Oxygen Tension at the Mitochondria (mmHg) | PmitO<sub>2</sub> | × | Γ(2, 8)<br>Min (0<br>mmHg)<br>Max (P<sub>cap</sub><br>mmHg) |
| Mean Transit Time through all vascular compartments (seconds) | MTT | 1.5 – 20 | 1.5 – 20 |
| Linear BOLD scaling term (s<sup>-1</sup>g<sup>-β</sup>dL<sup>β</sup>) | A | N (14, 2.0) | N (14, 2.0) |
| Non-linear BOLD scaling constant | β | 1.3 | 1.3 |
| Capillary Blood Volume (ml/100g) | CBV<sub>cap</sub> | 0.1 – 5 | 0.1 – 5 |
| Resting Cerebral Blood Flow (ml/100g/min) | CBF<sub>0</sub> | 15 – 180 | 15 – 180 |
| Arterial Blood Volume Fraction | - | 0.2 – 0.3 | 0.2 – 0.3 |
| Ratio between BOLD sensitive blood volume and capillary blood volume | ρ | 2 – 3.33 | 2 – 3.33 |
| Haemoglobin concentration (g/dL) | [Hb] | 14 | 14 |

When oxygen diffusivity is not included in the definition of M, but the physiology is constrained as per table 1, we are able to fit a Lowess surface (quadratic, span 25) to OEF_0_ (from M and CBF_0_) with an RMSE of 0.086 (R^2^ of 0.70) figure 1a. If oxygen diffusivity is included in the definition of M, figure 1b, we are able to fit a surface with an RMSE of 0.043 (R^2^ of 0.92), significantly reducing the error in OEF_0_. Thus, we are able to predict the modelled OEF_0_ with a RMSE error of approximately 10%, using only measures of M and CBF_0_ (given a fixed [Hb]).

**Figure 1.**
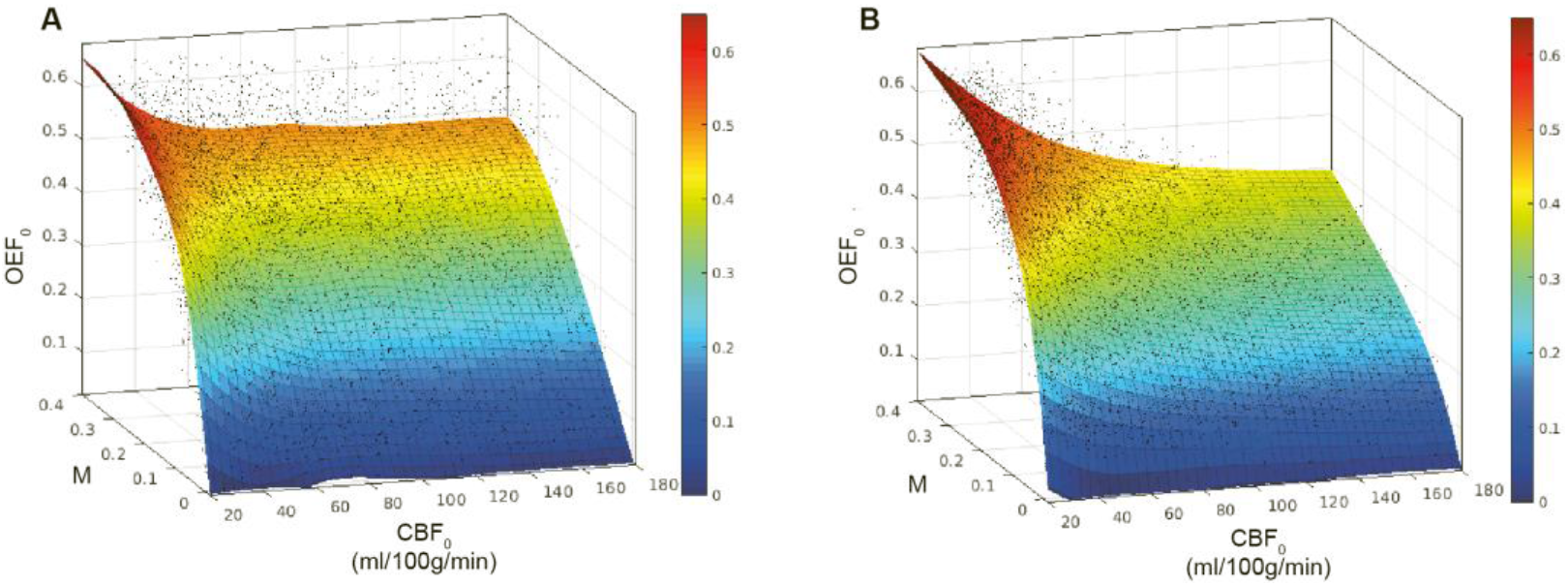
Lowess (quadratic) surface fits to modelled relationship between CBF_0_, M, and OEF_0_ with fixed [Hb]. **A**. Monte Carlo simulations with vascular volume fractions and MTT limited to physiological ranges (R^2^ = 0.70, RMSE 0.086). **B**. Monte Carlo simulations with proposed model of M including simple model of oxygen diffusion from capillary bed (R^2^ = 0.92, RMSE 0.043).

Because the Monte-Carlo simulations in figure 1 have a fixed [Hb] we cannot generalize the relationship across Hb concentrations. However, if a surface is fit for CMRO_2,0_ rather than OEF_0_, then [Hb] does not have to be considered as a separate parameter (at least under normoxic conditions). Figure 2 shows how a simple polynomial surface (order 2,3) can be fitted to M and CBF_0_ to estimate CMRO_2,0_ across a wide range of Hb values (10 to 18 g/dL) with a RMSE of 22.5 μmol/100g/min (R^2^ = 0.94).

**Figure 2.**
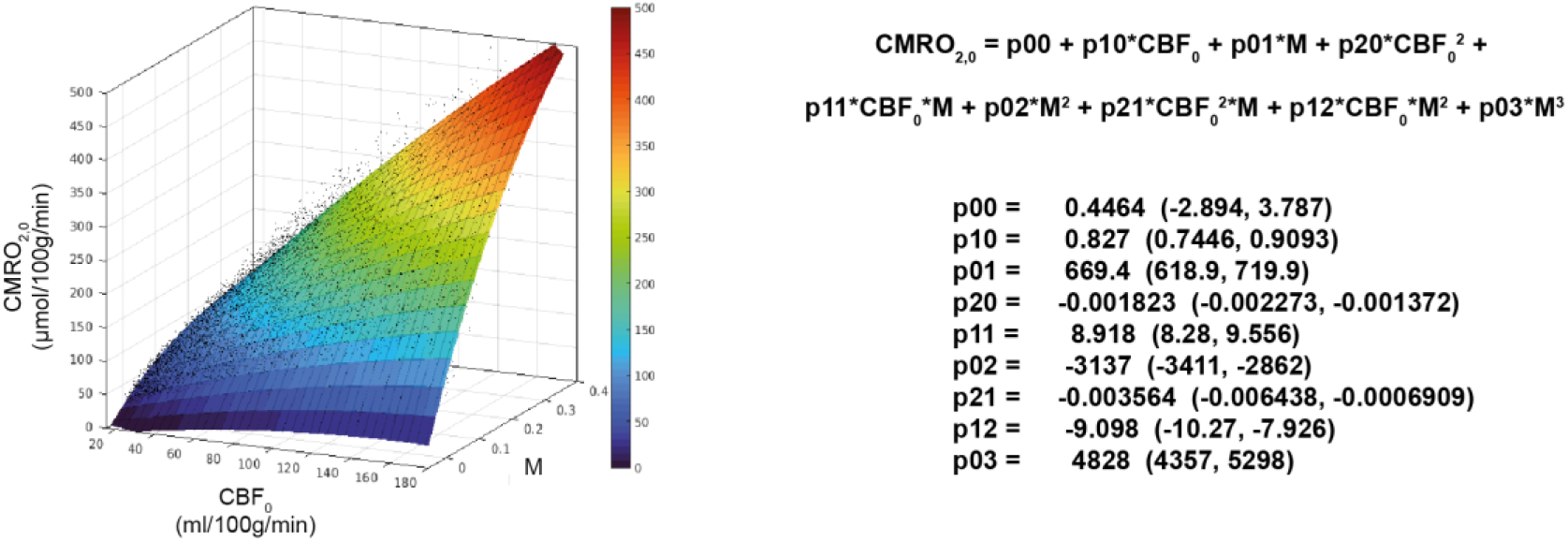
Polynomial surface fit to modelled relationship between CBF_0_, M, and CMRO_2,0_ with unknown [Hb]. Equation of the polynomial surface is presented with confidence intervals for each fitted parameter (R^2^ = 0.94, RMSE = 22.5 pmol/100g/min).

The inverse relationship, mapping CBF_0_ and CMRO_2,0_ to M, can also be estimated with a surface. Thus, allowing the maximum BOLD signal to be directly estimated from the physiological parameters. Equation 6 shows a rational equation that allows for such a mapping (RMSE = 0.04, R^2^ = 0.85).

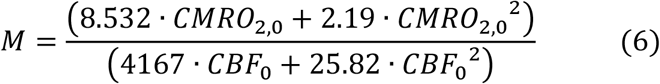

The modelling suggests that by fitting for M and CBF_0_, we can make estimates of CMRO_2,0_ with a low degree of uncertainty, across a wide range of physiology, and using only a single hypercapnic challenge. However, as highlighted by equation 3b, the BOLD response to breath-hold challenges has a dependence on CMRO_2,0_. Therefore, we cannot estimate CMRO_2,0_ in this step like manner (without making further approximations) and must instead make simultaneous estimates of M and CMRO_2,0_, for example, with numerical methods. Here we employ a gradient boosted decision tree algorithm, LightGBM, which is trained on simulated data to learn the mapping between MRI data and CMRO_2,0_. Thus, allowing rapid estimation of CMRO_2,0_ directly from the MRI data. The details of the machine learning analysis method and training are given in section 3.

## 3 Machine Learning Methodology

The machine learning regressors employed in this work are trained on simulated data. The method and its implementation are similar to our recently presented work on robust parameter estimation with dc-fMRI ^19^. In the present implementation the BOLD signal model (equations 3b and 5) and simplified pCASL kinetic arterial spin labelling signal model ^44^ were used to generate artificial MRI time series to match the in-vivo acquisition and breath-holding protocols.

The relaxation rate of arterial blood, R_1_, was modelled as in shown in equation 7 ^45, 46^ and the P50 for Hb saturation was estimated from the partial pressure of resting end-tidal CO_2_, as described in ^45^.

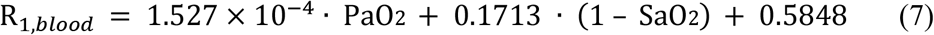

An acquisition length of 8 minutes 45 seconds was simulated with a TR of 4.4 seconds (119 volumes) and a BOLD echo time of 30ms. The breath-holding paradigm is detailed in figure 3, and table 2 lists the values and distributions of all MRI and physiological parameters used in the data simulations. Breath-holds were modelled using a boxcar function convolved with a gamma density function. Both hypercapnic and hypoxic variation was modelled, each with separate rise and fall times, taking into account the simultaneous changes in PaCO_2_ and PaO_2_ during breath-holding ^26^. Variation in local arrival times was modelled with a bulk delay of up to 3 TR’s (13.2 seconds).

**Table 2.** List of simulated MRI and physiological parameters used in ML data generation and their distributions.

| Parameter Description | Symbol | Range / Distribution |
| --- | --- | --- |
| Mean Transit Time through all vascular compartments (seconds) | MTT | 1.5 – 20 |
| Mean Capillary Transit Time (seconds) | $MTT_{cap}$ | 0.5 – 4.0 |
| Oxygen Tension at the Mitochondria (mmHg) | $P_{mitO_2}$ | $\Gamma(2, 8)$ |
| Linear BOLD scaling term ($s^{-1}g^{-\beta}dL^{\beta}$) | A | $N(14, 2.0)$ |
| Non-linear BOLD scaling constant | $\beta$ | 1.3 |
| Capillary Blood Volume (ml/100g) | $CBV_{cap}$ | 0.1 – 5 |
| Resting Cerebral Blood Flow (ml/100g/min) | $CBF_0$ | 0 – 200 |
| Arterial Blood Volume Fraction | - | 0.2 – 0.3 |
| Ratio between BOLD sensitive blood volume and capillary blood volume | $\rho$ | 2 – 3.33 |
| Oxygen Extraction Fraction | OEF | 0.05 – 0.65 |
| Haemoglobin concentration (g/dL) | [Hb] | 10 – 18 |
| Resting Arterial Oxygen Tension (mmHg) | $PaO_{2,0}$ | 90 – 130 |
| Change in Arterial Oxygen Tension (mmHg) | $\Delta PaO_2$ | -35 – -25 |
| Resting Arterial Carbon Dioxide Tension (mmHg) | $PaCO_{2,0}$ | 30 – 45 |
| Change in Arterial Carbon Dioxide Tension (mmHg) | $\Delta PaCO_2$ | 6 – 10 |
| Cerebral Vascular Reactivity (% CBF / mmHg) | CVR | 1 – 6 |
| Hemodynamic delay (seconds) | - | 0 – 13.2 |
| Shape of hemodynamic response function | - | $\Gamma(0.5 - 1, 1)$ |
| Effective permeability of capillary endothelium and brain tissue ($\mu\text{mol/mmHg/ml/min}$) | $\kappa$ | 3 |
| Hill Coefficient | h | 2.8 |
| Post Labelling Delay (seconds) | PLD | 1 – 3 |
| Tagging Duration (seconds) | - | 1.5 |

**Figure 3.**
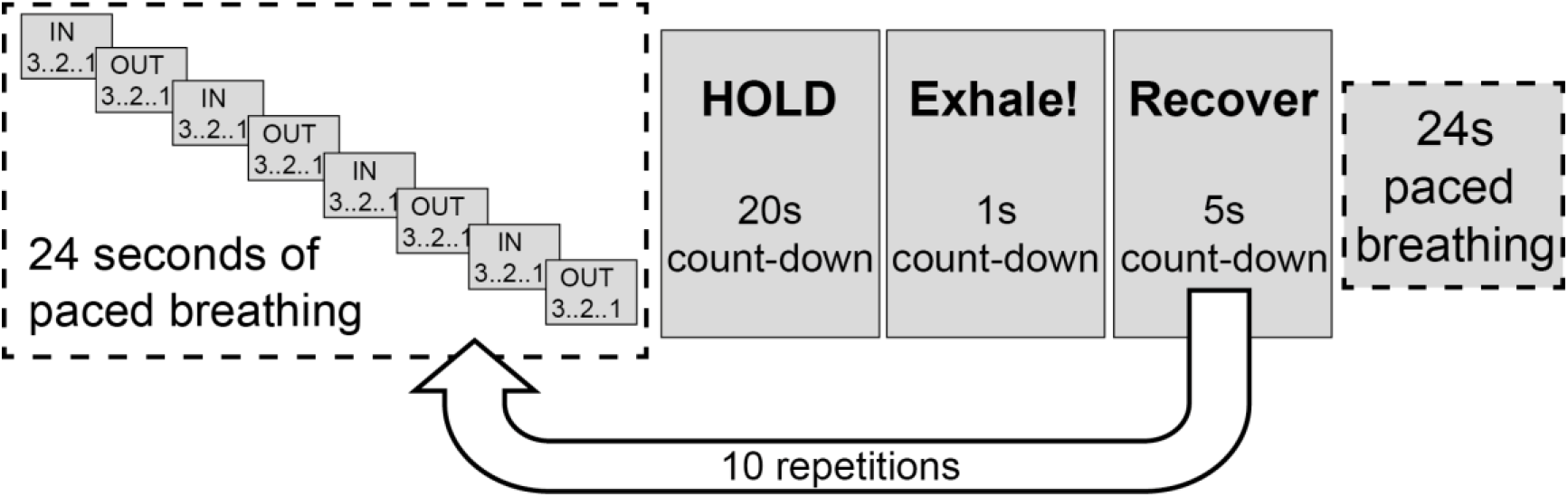
Breath-holding paradigm used to simulate machine learning data and for in-vivo data acquisition

Simulated MRI data was high pass filtered (300 second cut-off for the BOLD data only), and Fourier transformed to create frequency domain ASL and BOLD data. A feature vector was constructed by concatenating physiological and sequence parameters; [Hb], CaO_2,0_, post-label delay, and the first 15 points of the frequency domain data (excluding the DC BOLD component). The implementation in this work used only the MRI magnitude data (no phase data). This is a change to the previous implementation where both magnitude and phase data were used ^19^. The motivation for this change is to reduce the sensitivity to hemodynamic delays and is practicable due to the periodic nature of the breath-hold stimulus. We used LightGBM ^20^, an efficient gradient boosted decision tree, to learn the mapping between the MRI data and the two target parameters, CBF_0_ and CMRO_2,0_. LightGBM is a state-of-the-art gradient boosted decision tree algorithm that is computationally efficient and is suitable for training on big data sets. A summary of the machine-learning pipeline is presented in figure 4.

**Figure 4.**
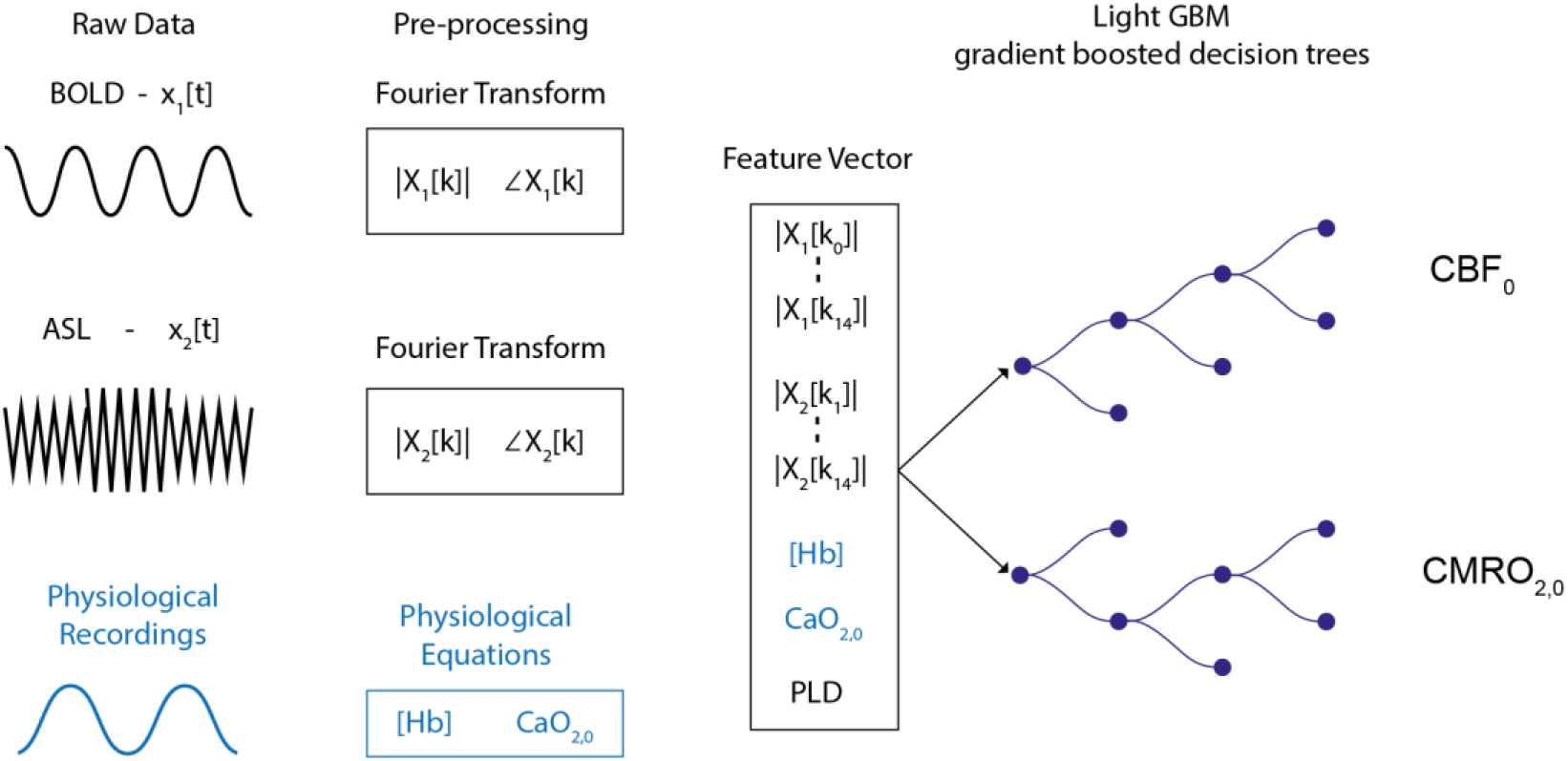
Schematic of the machine learning pipeline. Raw data is pre-processed and Fourier transformed to create a feature vector. LighGBM is used to train two independent regressor for estimating baseline perfusion and the resting cerebral metabolic rate of oxygen consumption.

50,000 simulations were used for training an individual regressor to predict CBF_0_ and CMRO_2,0_. Training of each network took less than 10 seconds on a 2013 MacBook Pro with 16GB of memory. We used the Python implementation of LightGBM (version 3.0.0.99) for training with the maximum number of leaves per tree set to 1,000; all other parameters were default. To assess the performance of the LightGBM implementation 10,000 additional simulations were performed using the distribution of parameters detailed in table 2, but with OEF_0_ limited to 0.15 to 0.65.

## 4 In-vivo experiments

Fifteen healthy volunteers (4 males, mean age 27.4 ± 6.2 years) were recruited to the study. The local ethics committee approved the study and written informed consent was obtained from each participant. Blood samples were drawn via a finger prick prior to scanning and were analysed with the HemoCue Hb 301 System (HemoCue, Ängelholm, Sweden) to calculate the systemic [Hb] value for each participant. All MRI data were acquired using a Siemens MAGNETOM Prisma (Siemens Healthcare GmbH, Erlangen) 3T clinical scanner with a 32-channel receiver head coil (Siemens Healthcare GmbH, Erlangen). An 8 minute 45 second dual-excitation pseudo-continuous arterial spin labelling (pCASL) and BOLD-weighted acquisition was acquired during repeated breath-holding (see figure 3 for experimental protocol). End-tidal gas monitoring was performed throughout the acquisition via a nasal cannula using a rapidly responding gas analyser (PowerLab®, ADInstruments, Sydney, Australia). All volunteers practised the breath-hold task to ensure they understood the visual instructions, prior to the MRI session.

The parameters for the in-house pCASL sequence ^45^ were as follows: post-labelling delay and label duration 1.5 seconds, EPI readout with GRAPPA acceleration (factor = 3), TE_1_ = 10ms, TE_2_ = 30ms, TR = 4.4 seconds, 3.4 x 3.4 mm in-plane resolution, and 15 (7mm) slices with 20% slice gap. A 3D magnetization-prepared rapid gradient-echo (MPRAGE) sequence was acquired for image registration purposes (1mm slice thickness, 1.14 × 1.14 in-plane image resolution, TR/TE = 2100/3.2 ms). FAST ^47^ was used for tissue segmentation to create high-resolution grey matter partial volume estimates.

Post processing of the pCASL data included motion correction of the time series with the FSL tool MCFLIRT ^48^ followed by surround subtraction and surround averaging of TE_1_ and TE_2_ time series to create perfusion-weighted and BOLD-weighted time series respectively. After temporal smoothing, the BOLD time series was high pass filtered with a 300 second cut-off using the filter implementation in FSL. Each time series was then Fourier transformed, and the first 15 points of the magnitude data were included in a feature vector used for parameter estimation. The DC component from the BOLD data was excluded from the feature vector such that only 14 BOLD data points were included. The feature vector was completed with the inclusion of CaO_2,0_, [Hb], and the post-labelling delay (calculated per slice). CBF_0_ and CMRO_2,0_ parameter estimation was performed using the LightGBM estimator trained on simulated data.

Subjects’ resting perfusion estimates were used to register the low-resolution functional data to high-resolution grey matter partial volume estimates. An inverse transform was used to create partial volume estimates in functional space. A grey matter partial volume estimate of 0.5 was taken as the threshold to create grey matter masks for statistical reporting of grey matter averages.

## 5 Results - Simulations

Cross-validation of the regression models show that CBF_0_ estimates have an R^2^ of 0.99 and a RMSE of 0.3 ml/100g/min, while CMRO_2,0_ estimates have an R^2^ of 0.94 and a RMSE of 22.9 μmol/100g/min. This is a similar level of uncertainty observed in section 2, when using known M and CBF_0_ values, demonstrating the ability of the regressor to estimate CMRO_2,0_ directly from the simulated MRI data. In addition to quantifying the RMSE for individual voxels it is equally important to assess the bias in parameter estimates and their sensitivity to variation in cerebral physiology. Figure 5 shows the percentage error in CMRO_2,0_ estimates for key modelling parameters. The error plots typically have a bias of less than 10%, with maximum values approaching 20% at the extreme ranges of parameter values.

**Figure 5.**
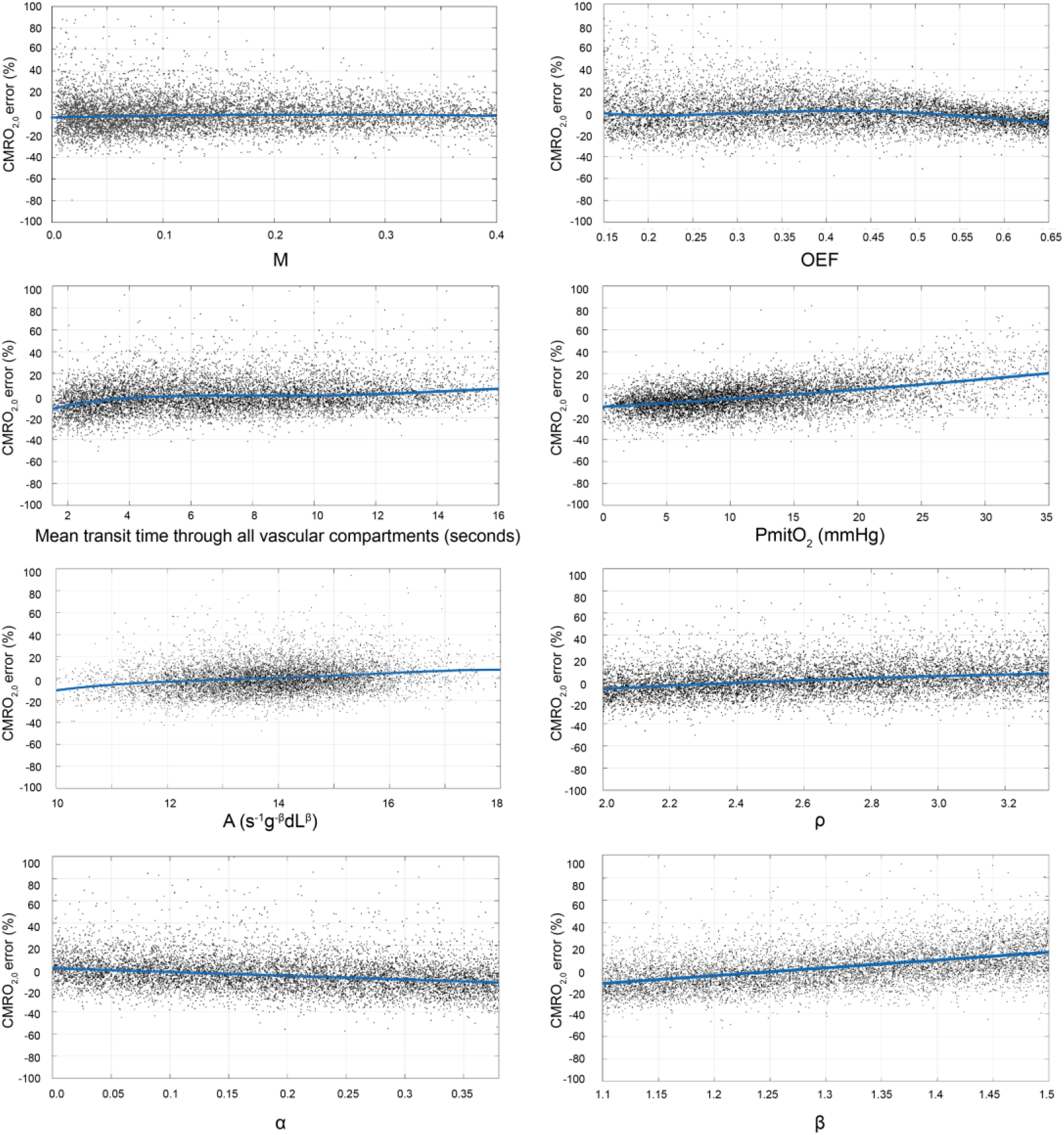
Monte-Carlo simulation to assess the error and bias in CMR0_2,0_ estimates with respect to modelled physiological parameters. Each plot displays the percentage error in CMRO_2,0_ estimates made with the trained LightGBM regressor on simulated data. The lines of best fit are nth order polynomials where n was chosen to minimise the adjusted RMSE in each plot.

The modelling parameter with the largest bias is PmitO_2_. However, if we plot a surface of the mean percentage CMRO_2,0_ error against MTT and PmitO_2_, figure 6, we see that the most significant bias only occurs when MTT is long and PmitO_2_ is high. Although this is potentially problematic in certain pathologies, it is unlikely to present a practical limitation, as ASL imaging at 3T is unlikely to provide useful data when transit times are very long.

**Figure 6.**
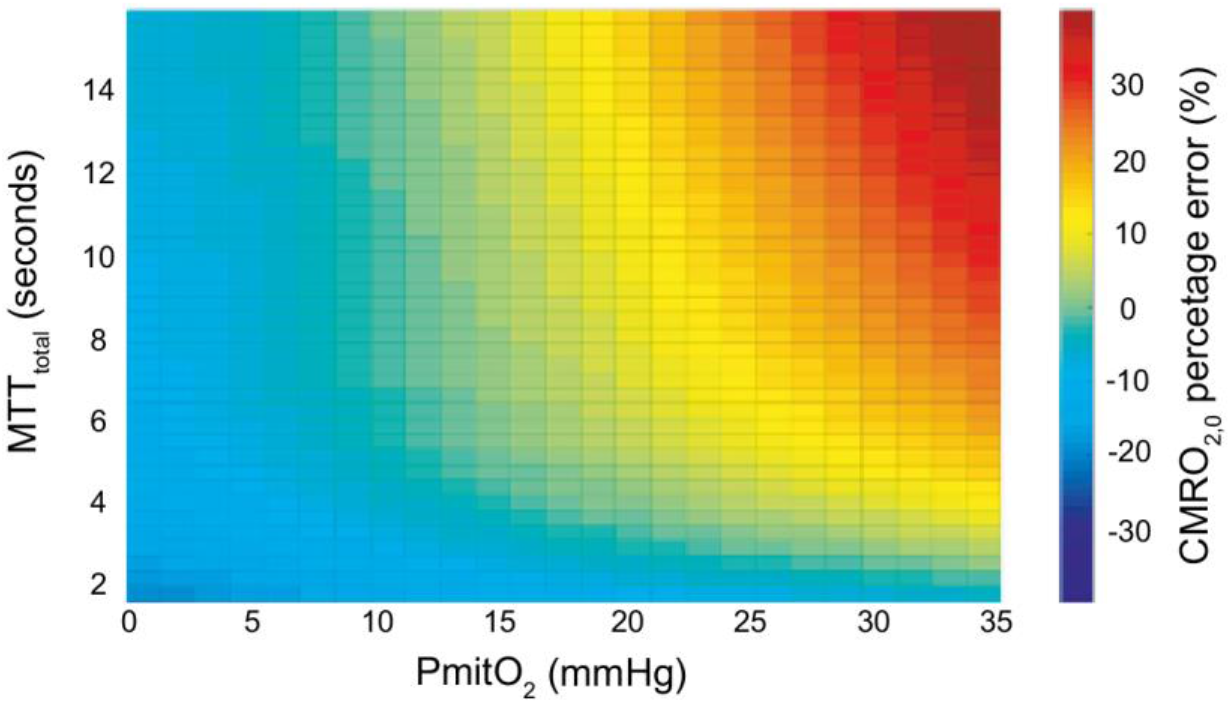
Mean percentage error in CMRO_2,0_ plotted against PmitO_2_ (mmHg) and total MTT (seconds). The error in CMRO_2,0_ is small over most of the physiological range, only increasing significantly when both MTT is long and PmitO_2_ is high

Taken together, these results demonstrate that the implemented ML algorithm is able to accurately capture the modelled relationship between the MRI data and the physiological parameters across a wide distribution of physiological parameters. The lack of significant biases suggests the method should be sensitive to metabolic variation in both healthy and pathological tissue when applied in-vivo.

## 5 Results - In Vivo

All subjects were able to follow the experimental instructions and successfully performed the respiratory task. We were unable to obtain a blood sample from one subject out of fifteen scanned. This subject was excluded from further analysis, as they did not have a measured [Hb]. The average grey matter temporal signal (tSNR) to noise was 2.8 for the ASL acquisitions and 90 for the BOLD acquisitions.

Table 3 provides a summary of average grey matter results each for subject. M values were calculated voxelwise using equation 6 to find an M value consistent with the estimated CBF_0_ and CMRO_2,0_. Matching parameter maps from each subject are displayed in figure 7.

**Table 3.** Mean grey matter parameter estimates and systemic [Hb] measurements for each subject.

| Subject | [Hb]<br>(g/dL) | OEF | CBF<br>(ml/100g/min) | $\text{CMRO}_2$<br>($\mu\text{mol}/100\text{g}/\text{min}$) | M<br>(%) |
| --- | --- | --- | --- | --- | --- |
| 1 | 14.6 | 0.43 | 35.7 | 110.1 | 15.9 |
| 2 | 10.4 | 0.48 | 42.9 | 108.1 | 12.3 |
| 3 | 16.1 | 0.44 | 41.2 | 144.3 | 22.3 |
| 4 | 13.0 | 0.42 | 45.3 | 123.9 | 15.2 |
| 5 | 13.0 | 0.43 | 42.1 | 118.8 | 15.1 |
| 6 | 13.4 | 0.44 | 40.1 | 121.9 | 16.4 |
| 7 | 18.5 | 0.36 | 37.2 | 117.3 | 17.6 |
| 8 | 13.6 | 0.39 | 47.7 | 126.8 | 14.9 |
| 9 | 12.7 | 0.40 | 59.5 | 153.2 | 16.5 |
| 10 | 13.0 | 0.43 | 46.4 | 132.6 | 16.5 |
| 11 | 14.7 | 0.39 | 49.6 | 146.4 | 18.9 |
| 12 | 13.4 | 0.41 | 39.4 | 108.9 | 13.9 |
| 13 | 14.2 | 0.40 | 49.2 | 140.5 | 17.6 |
| 14 | 13.9 | 0.39 | 42.6 | 116.4 | 14.4 |
| **Mean** | **13.9** | **0.42** | **44.2** | **126.4** | **16.3** |
| **Std Dev** | 1.8 | 0.03 | 6.0 | 14.3 | 2.3 |
| **COV** | 0.13 | 0.07 | 0.13 | 0.11 | 0.14 |

**Figure 7.**
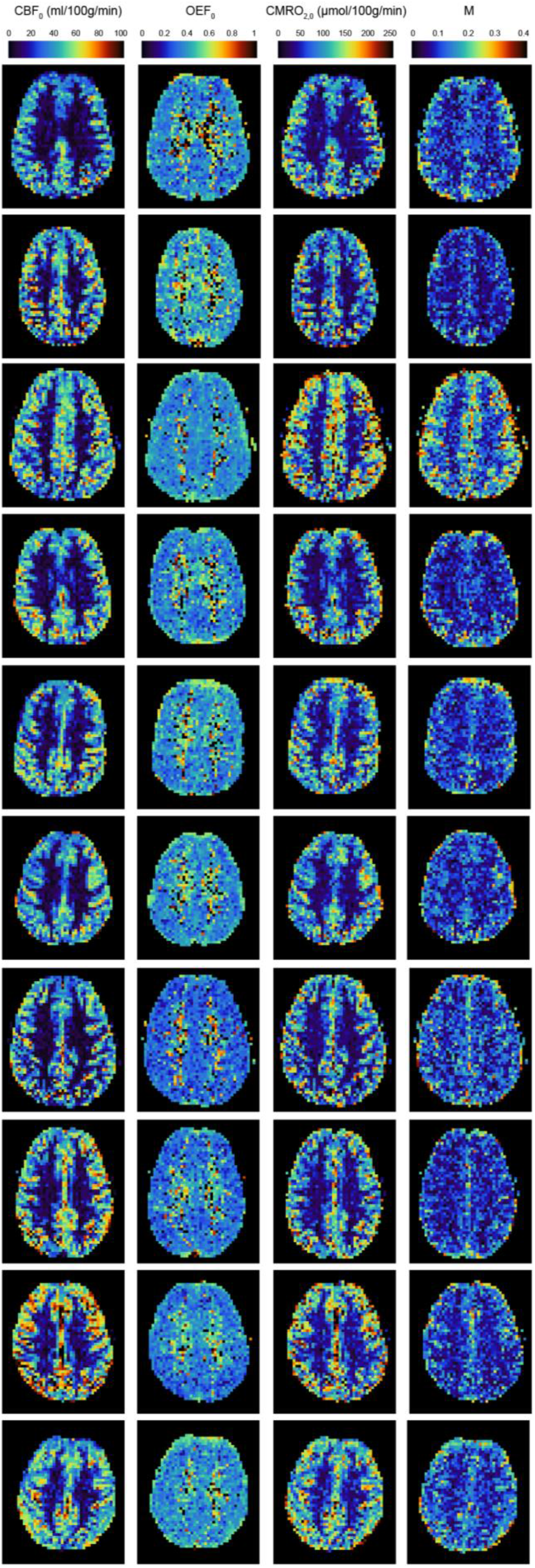
Example parameter maps for the first 10 subjects scanned. A single slice was chosen to display CBF_0_, OEF_0_. CMRO_2,0_, and M parameter estimates. OEF_0_ estimates are approximately unifrom within grey matter. However, OEF_0_ is elevated in white matter. Overestimation in white matter is likely due to long blood arrival times causing ASLto underestimate perfusion changes during breath-holding.

Figure 7 shows example CBF_0_, OEF_0_, CMRO_2,0_, and M maps for each subject. As can be seen from the figure, and consistent with expected physiology, regions of high perfusion are matched with areas of high metabolic oxygen consumption, while the oxygen extraction fraction has little variation throughout the grey matter. However, OEF_0_ estimates in pure white matter voxels are artificially high. This is likely to be a consequence of the long arrival time of blood in white matter leading to a reduced perfusion signal. In accordance with expectations, the M maps contain higher values in grey matter compared to white matter (due to increased blood volume). However, due to the overestimation of OEF_0_ in white matter M is also overestimated here, reducing the expected grey matter to white matter contrast.

A regression of grey matter OEF percentage against [Hb] and CBF produced the following model (p < 0.05).

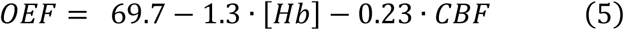

The parameter fits are significant for both CBF and [Hb] (p<0.05) and are in close agreement with the results reported by Ibaraki et al 2010 ^49^ and with our previous work using dual-calibrated fMRI. This result suggests that the magnitude of parameter estimates made with the proposed method are appropriate, and that the method is able to detect underlying variations in physiology and metabolism, as predicted by the modelling.

## 5 Discussion

This work presents a practical and pragmatic method for mapping the rate of cerebral grey matter metabolic oxygen consumption with MRI. The approach is straightforward to apply and should be widely applicable to research as well as clinical studies. Although the approach requires validation with proven measures of cerebral oxygen metabolism, the agreement between in-vivo results and previously observed physiological relationships is encouraging and supports the validity of the method.

We have chosen to demonstrate the method using a breath-holding protocol with simultaneously acquired BOLD and ASL data. This method provides rapid measurement of both M and CBF_0_, which are required for the calculation of oxygen extraction and CMRO_2_. We have also implemented a machine-learning analysis approach for rapid parameter estimation. However, the method can in principle be applied to any data that can provide a robust measurement of M and an estimate of resting CBF. To facilitate such analysis, we have provided a simple mapping from M and CBF_0_ to CMRO_2,0_.

We show robust parameter estimates in a healthy young cohort based on a breath-hold of 20 seconds, repeated 10 times, and interleaved with paced breathing. Many breath-hold repeats are needed to ensure sufficient signal to noise when modelling ASL data with a long effective TR. Despite some extra challenges that may arise for clinical applications, this breath-hold method is non-invasive and does not require specialist respiratory equipment for gas inhalation, therefore it has potential for wider applicability in research and clinical settings. Breath holding tasks during MRI have been used successfully in many clinical populations ^50–56^, mostly during BOLD fMRI to map cerebrovascular reactivity. Task cues and practice sessions may need to be tailored to each clinical population to ensure adequate compliance.

The limitations of the method are mostly shared with the dual-calibrated methodology, see ^57^ for a review. For example, for the method to be viable there must be a local increase in CBF with breath-holding, i.e., there must be a local vascular reserve. This condition may not be met in diseases such ischemic stroke, where arterial vessels are maximally dilated in an attempt to maintain local perfusion. Additionally, the method is expected to be vulnerable to large changes in the ratio of the capillary to deoxyhaemoglobin sensitive blood volumes, i.e. beyond the typical variation in vascular compartments referenced in this work. Although large changes in this ratio appear unlikely in many brain diseases, we might expect such vascular remodelling to occur in advanced or aggressive brain tumours ^58^. Therefore, we would encourage caution in interpreting the results in such applications. Nevertheless, the proposed method offers a simple means of mapping cerebral oxygen metabolism with MRI and has the potential to be a useful tool for both neuroscience research and clinical imaging.

## Acknowledgements

We would like to thank the UK Engineering and Physical Sciences Research Council (EP/S025901/1) and Wellcome for supporting this work: Wellcome Strategic Award, Multi-scale and multi-modal assessment of coupling in the healthy and diseased brain, grant reference 104943/Z/14/Z. MG thanks the Wellcome Trust for its generous support via a Sir Henry Dale Fellowship (220575/Z/20/Z).

## Data Availability Statement

The python code for the data analysis pipeline proposed in this article is freely available https://zenodo.org/badge/latestdoi/350795130. We do not have ethical consent to make the in-vivo datasets acquired for this study publicly available.

